# Shining a light on the dark Nt-acetylome by integrating omics data

**DOI:** 10.64898/2025.12.31.697169

**Authors:** Salomé Nashed, Médine Benchouaia, Angélie Dijoux-Maréchal, Thierry Delaveau, Stéphane Le Crom, Benoit Palancade, Frédéric Devaux, Mathilde Garcia

## Abstract

N-terminal acetylation, catalysed by N-terminal acetyltransferases (Nats), is one of the most prevalent protein modifications and is implicated in human diseases. Yet, despite extensive COFRADIC-based proteomics, only ∼5–10% of the proteome has been interrogated, leaving the majority of the Nt-acetylome unexplored. Here, we combined all major COFRADIC datasets with sel-TRAP, a high-sensitivity, orthogonal approach for profiling co-translational Nat targets via selective ribosome purification. This integrated analysis refined the substrate specificities of NatA, NatB, and NatC/E/F and provided the most comprehensive view to date of the canonical human and yeast Nt-acetylomes. Importantly, we also uncovered hundreds of cryptic Nat substrates arising from alternative translation initiation, with unexpected Nt-proteoforms constituting a previously underappreciated source of Nat targets. Collectively, our results revealed the complex landscape of a “dark” Nt-acetylome, the characterization of which, including its functional roles in regulating protein function and in disease, remains a major challenge for future research.

## INTRODUCTION

Protein N-terminal (Nt) acetylation (NTA) is a common protein maturation process that occurs mainly co-translationally, as the nascent polypeptide emerges from the ribosome (for an overview of NTA mechanism and biological function see review^1^). This irreversible modification, which neutralizes the positive charge of the N-terminal amine and alters its hydrophobicity, is catalyzed by specific N-terminal acetyltransferases (Nats). Among the Nat complexes, NatA, NatB, and NatC are conserved across all eukaryotes, whereas NatE is present but inactive in yeasts. Notably, additional NATs, NatF–H, are absent in yeasts and present in metazoans and plants, where they can act post-translationally, expanding the scope of this modification beyond co-translational processing. NTA has been estimated to affect up to 80% of human and plant proteins and about 60% of yeast proteins^1,2^. It can influence multiple aspects of protein biology, including folding, protein complex formation, subcellular targeting, and degradation, as shown in various model systems^1^. The pathological relevance of NTA is underscored by its implication in diverse human diseases such as cancer, developmental syndromes, and neurodegeneration diseases, as well as in physiological defects in plants^1^.

Recent advances in Nt proteomic techniques have enabled the capture of thousands of cellular Nt peptides and the quantification of their acetylation levels. These studies have generated quantitative maps of NTA levels for the most highly expressed proteins across multiple cellular models, including *S. cerevisiae*, *H. sapiens*, and *A. thaliana*^2–13^, thereby laying the groundwork for a systematic understanding of Nt-acetylation. Among the available approaches, Nt-COFRADIC (combined fractional diagonal chromatography) remains the gold standard for analyzing *in vivo* Nt-acetylomes (for an overview of Nt-acetylation experimental techniques, see review^2^). This technique primarily relies on the mass spectrometric analysis of purified N-terminal peptides. The percentage of their *in vivo* acetylation is estimated by assessing their light/heavy ratio following *in vitro* acetylation of the free N-termini with heavy d(3)-acetyl^3^. This method has been used to identify yeast and human Nats substrates by examining changes in acetylation stoichiometry of N-termini in knockout or knockdown experiments, as well as through ectopic expression of human Nats in yeast cells^4–8^ (see Table 1). These data, recapitulated in Supplemental Figures 1 and 2, confirmed the canonical substrate specificities described in the previous literature: NatA targets small residues (Ser, Ala, Thr, Gly, Val) at the N-terminus following removal of the initiator methionine (iMet) by methionine aminopeptidases (MetAP); NatB and NatC directly acetylate retained iMet when followed by specific residues, i.e. Asn/Asp or Gln/Glu for NatB, and large hydrophobic residues (Phe, Leu, Met, Trp, Ile, Tyr) for NatC. These data also indicated that MH-N-termini may be extended substrates of NatB^4^, while MK- N-termini may be extended substrates of NatC^9^, although these conclusions are based on a limited number of identified N-terminal peptides. Additionally, NatC has been found responsible for the NTA of MS- and MV- N-termini resulting from atypical retention of iMet^9^.

**Table 1.**
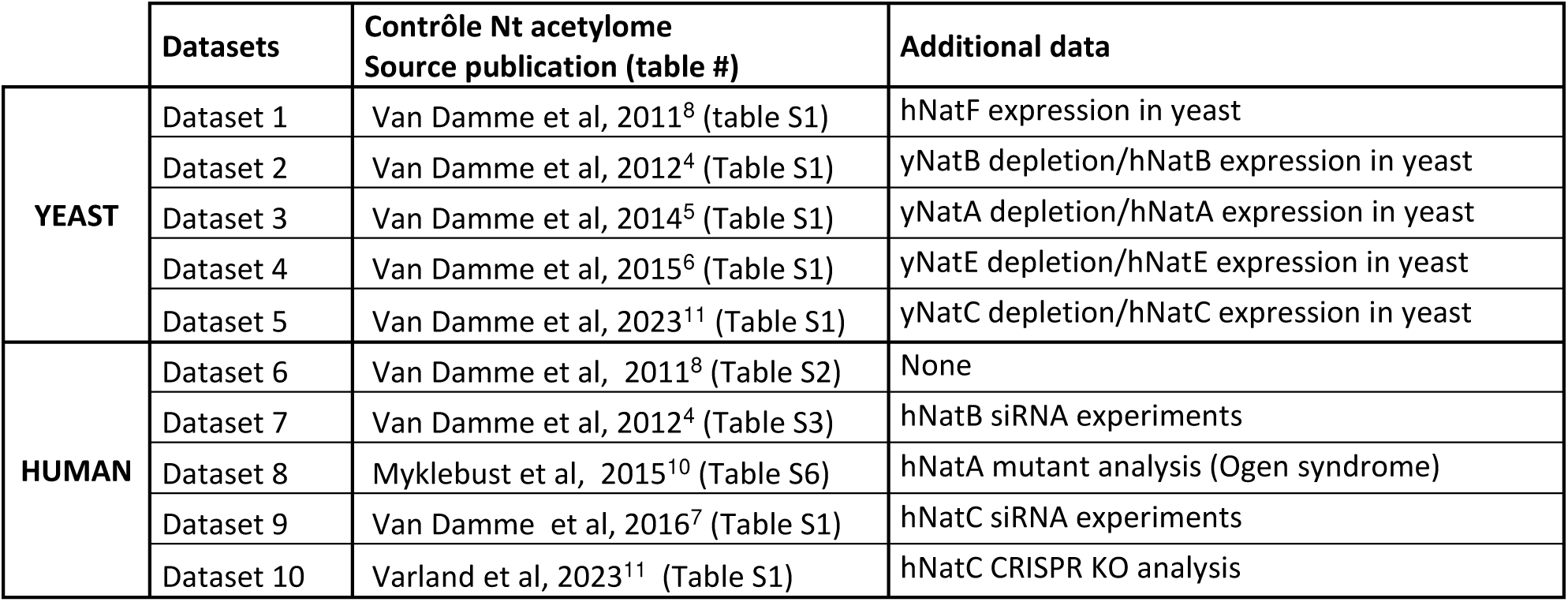

|  | Datasets | Contrôle Nt acetylome<br>Source publication (table #) | Additional data |
| --- | --- | --- | --- |
| **YEAST** | Dataset 1 | Van Damme et al, 2011<sup>8</sup> (table S1) | hNatF expression in yeast |
|  | Dataset 2 | Van Damme et al, 2012<sup>4</sup> (Table S1) | yNatB depletion/hNatB expression in yeast |
|  | Dataset 3 | Van Damme et al, 2014<sup>5</sup> (Table S1) | yNatA depletion/hNatA expression in yeast |
|  | Dataset 4 | Van Damme et al, 2015<sup>6</sup> (Table S1) | yNatE depletion/hNatE expression in yeast |
|  | Dataset 5 | Van Damme et al, 2023<sup>11</sup> (Table S1) | yNatC depletion/hNatC expression in yeast |
| **HUMAN** | Dataset 6 | Van Damme et al, 2011<sup>8</sup> (Table S2) | None |
|  | Dataset 7 | Van Damme et al, 2012<sup>4</sup> (Table S3) | hNatB siRNA experiments |
|  | Dataset 8 | Myklebust et al, 2015<sup>10</sup> (Table S6) | hNatA mutant analysis (Ogen syndrome) |
|  | Dataset 9 | Van Damme et al, 2016<sup>7</sup> (Table S1) | hNatC siRNA experiments |
|  | Dataset 10 | Varland et al, 2023<sup>11</sup> (Table S1) | hNatC CRISPR KO analysis |

While these studies established key principles of Nat substrate specificity, they were limited by the number of identified N-terminal peptides, typically representing only 5–10% of the analysed proteome. To overcome this limitation and expand our understanding of the Nt-acetylome, we integrated multiple COFRADIC datasets in yeast and humans^4–11^ (Table 1) and complemented them with sel-TRAP (Selective Translating Ribosome Affinity Purification) analyses in yeast, enabling sensitive transcriptomic detection of Nat co-translational targets from co-captured mRNAs^14^. Statistical analyses of the combined datasets refined NatA, NatB, and NatC substrate preferences, confirming canonical specificities and revealing newly extended ones, which allowed us to assemble the most comprehensive inventory of Nat substrates to date (1,145 in yeast and 1,683 in humans).

Importantly, our analyses also reveal a “dark” Nt-acetylome: many Nat substrates remain missing from current datasets and cannot be reliably predicted due to multiple phenomena that we quantified and exemplified. For example, unconventional iMet acetylation and alternative Nt-proteoforms, differing in N-terminal residues, iMet removal, and Nt-acetylation status, substantially diversify the Nt-acetylation landscape. These variations generate atypical N-termini that have been largely overlooked in previous analyses. By integrating proteomic and transcriptomic datasets, we notably identified 104 yeast and 123 human cryptic Nat substrates produced by alternative translation initiation. These alternative start substrates may modulate proteins with multiple cellular localizations, as exemplified by yeast fumarase, whose mitochondrial and nuclear activities could be regulated by distinct Nt-proteoforms. Overall, our work highlights the conceptual challenge of capturing the full complexity of Nt-proteoforms and integrating this knowledge into models of cellular physiology and human disease

## RESULTS AND DISCUSSION

### Meta-analysis of N-terminal proteomic datasets: quantitative insights into Nat specificities

Between 2011 and 2023, five Nt-COFRADIC datasets have been published for yeast and human cell lines (Table 1, Supplementary data file 1, Figure 1). These datasets cover only a fraction of the canonical annotated N-termini (peptides initiating at positions 1 or 2 of the annotated CDSs, depending on iMet removal) of the yeast proteome (7-13%) and the human proteome (4-6%) (Figure 1A and B, left panels, Table S1). Combining these data for a meta-analysis of the yeast and human Nt-acetylome resulted in an increase in proteome coverage of up to 19% for yeast and 9% for human, with acetylation measurements (% Nt acetylation) for 1,172 and 1,931 N-termini respectively (Figure 1A and B, left panels and Table S1). The number and coverage of N-termini, classified according to the residue following the initiator methionine (iMet), vary considerably depending on the amino acid considered (Figures 1A and B, right panels). This uneven representation introduces heterogeneity, which can significantly affect the statistical reliability of extrapolating the observed N-terminal acetylation frequencies to the entire proteome. Notably, in both yeast and human datasets, NatC-type N-termini with large hydrophobic residues (MF-, ML-, MI-, MW-, MM-, MY-) are each represented by fewer than 50 peptides, resulting in low statistical power for predicting their acetylation frequencies at proteome level (see confident intervals in Figures 2 A3 and B3, histograms).

**Fig 1.**
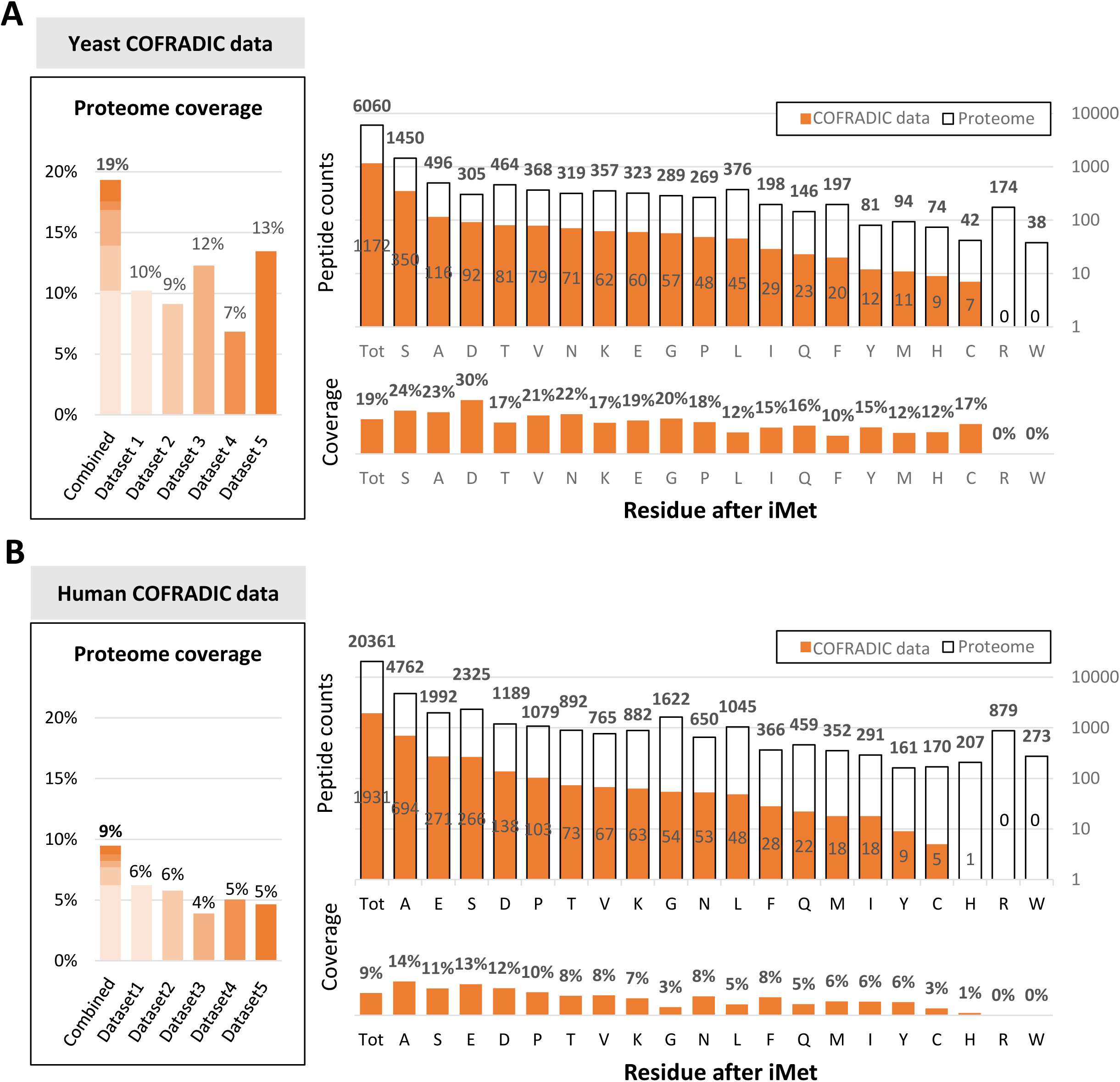
Increased Nt-acetylome coverage via dataset integration For yeast (A) and humans (B), five COFRADIC datasets were combined (Table 1), covering 7–13% and 4–6% of annotated canonical N-termini, respectively, thereby extending overall proteome detection to 19% (1,172/6,060) in yeast and 9% (1,931/20,361) in humans (left bar charts). Residues following iMet showed heterogeneous representation in the combined datasets (0–30% in yeast, 0–14% in humans), with some N-termini type observed more than 100 times and others fewer than 10 times (right bar charts). Peptide counts and detection frequencies are shown in the top and bottom right bar charts, respectively; orange bars indicate N-termini detected by COFRADIC, and white bars indicate all annotated N-termini, with the number of peptides shown in the corresponding areas.

**Fig 2.**
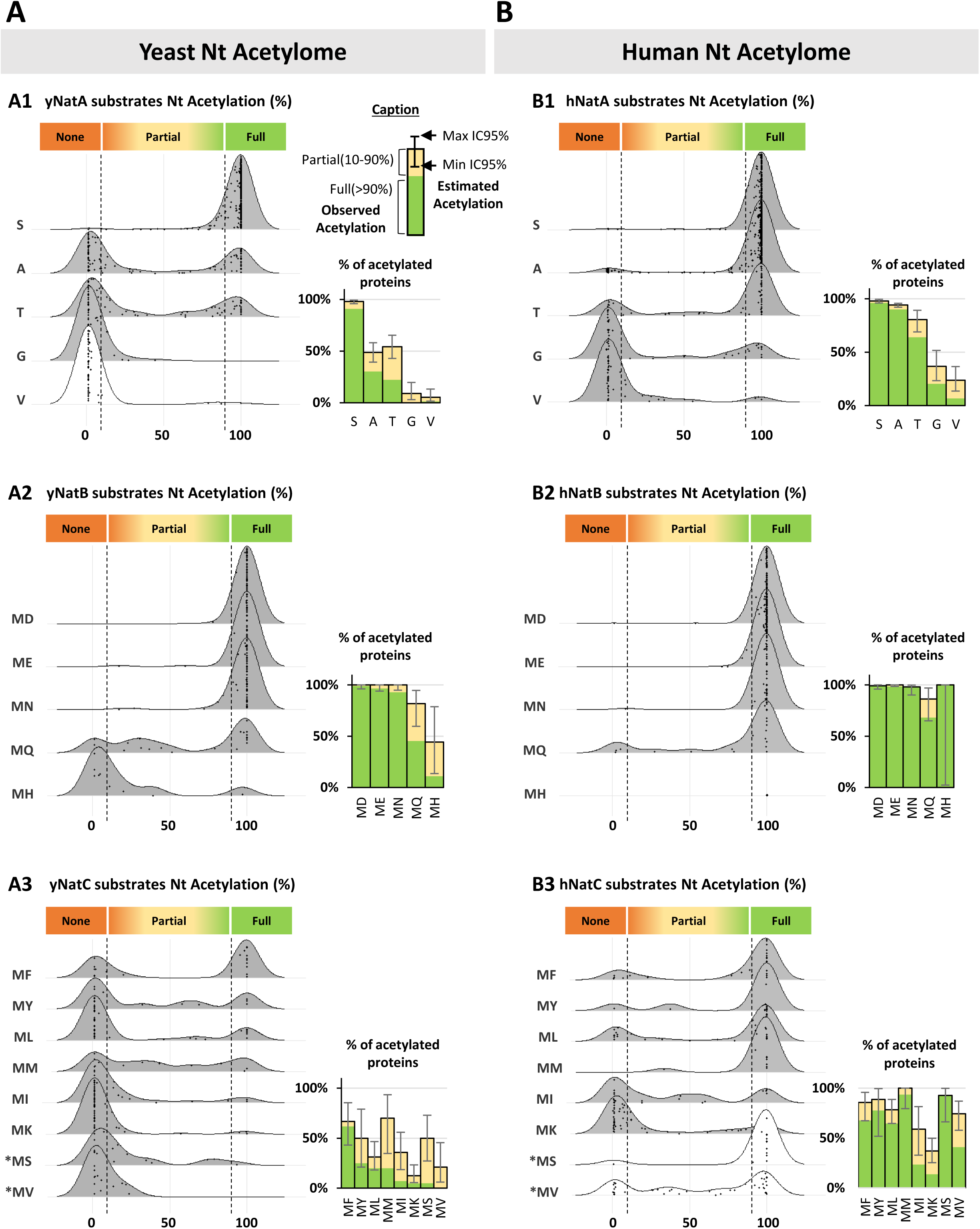
Overview of the human and yeast Nt-acetylome from combined datasets Yeast (A) and human (B) N-terminal peptides were classified as NatA (A1, B1), NatB (A2, B2), or NatC (A3, B3) putative substrates, and their Nt-acetylation percentage distributions are shown as ridgeline plots (Left), with individual N-terminal peptides represented as dots. Substrate specificity (NatA: S, A, T, G, V; NatB: MD, MN, MQ, ME, MH; NatC: MF, MY, ML, MM, MI, MK, MS, MV) was defined from yeast Nat deletions and human Nat overexpression in Nat-deleted strains (Supplementary Figs. 1–2). White plots (instead of grey) indicate cases where specificity could not be confirmed in one species (human or yeast) and was inferred from the other. Peptides were further classified as not acetylated (NTA < 10%), partially acetylated (10–90%), or fully acetylated (NTA > 90%). Right-hand histograms show the percentage of acetylated peptides (green: fully acetylated, yellow: partially acetylated), with 5% confidence intervals (exact binomial test) for extrapolation to the full proteome.

From these combined datasets, we addressed the challenge of integrative quantitative visualization of Nat substrate specificity using ridgeline plots (Figure 2A–B). This representation highlights both the global acetylation distribution of each substrate class and the acetylation levels of individual experimentally detected peptides, which we classified according to their acetylation status: (1) non-acetylated (NTA < 10%), (2) partially acetylated (10% < NTA < 90%), and (3) fully acetylated (NTA > 90%). We complemented this visualization with statistical analyses quantifying, for each substrate class, the fraction of acetylated peptides. To do so, we computed in both species the 5% confidence interval of acetylation rates for each N-terminal type, providing an estimate of the predictive capacity of current datasets for inferring the cellular Nt-acetylome (Figure 2A–B, histograms).

Confidence in extrapolating NTA depends on the Nat, the N-terminus, and the species considered. Some N-termini are well represented and consistently acetylated, allowing high-confidence extrapolations. This is the case in both yeast and human for S- N-termini targeted by NatA (Figures 2 A1 and B1, histograms) and most typical NatB targets (MD-, ME-, MN-) (Figures 2 A2 and B2, histograms). Other NatA substrates (A-, T-, G-, V-) and NatB substrates (MQ-) do not reach systematic acetylation particularly in yeast, making the acetylation of undetected similar N-termini less predictable. For instance, in yeast, only ∼50% of the detected proteins with A- or T-type N-termini are acetylated, and very few with G- or V-type N-termini (Figure 2 A1, histogram). Partial acetylation further complicates NTA extrapolation, as both acetylated and non-acetylated protein forms coexist in the cell in proportions that cannot be accurately predicted. Finally, low coverage of NatC substrates limits reliable predictions in both species, as reflected by wide confidence intervals, indicating that the detected samples may not be fully representative (Figures 2 A3 and B3, histograms).

Overall, human protein NTA is more predictable than in yeast, due to generally higher acetylation levels on the detected proteins (Figure 2). For example, NatA substrates, particularly A- and T-type N-termini (Figure 2 A1 and B1), show higher acetylation in human than in yeast, resulting in more reliable NTA predictions. Since NatA is the only enzyme capable of acetylating N-terminal residues after iMet removal^1^, this increase cannot be attributed to additional Nats in human cells compared to yeast, and likely reflects greater efficiency of human NatA. We also observed higher iMet acetylation in human, including acetylation of non-canonical N-termini resulting from iMet removal failure (i.e., MS- and MV-type N-termini; Figures 2A3 and 2B3). This is likely due to the presence of additional Nats in metazoans, namely NatE and NatF^1^. Indeed, data from ectopic expression of human NatE and NatF in yeast^6,8^ showed that these enzymes can extend iMet NTA by acetylating N-termini that escape modification by yeast NatB or NatC (Figure 3A). This includes NatC substrates with large hydrophobic residues after iMet, as well as MQ-type N-termini that are not consistently acetylated by NatB. In addition, NatE and NatF, like NatC, can acetylate iMet resulting from iMet removal failure (i.e., MT-, MS-, MA-, MV-type N-termini; Figure 3A).

**Fig 3.**
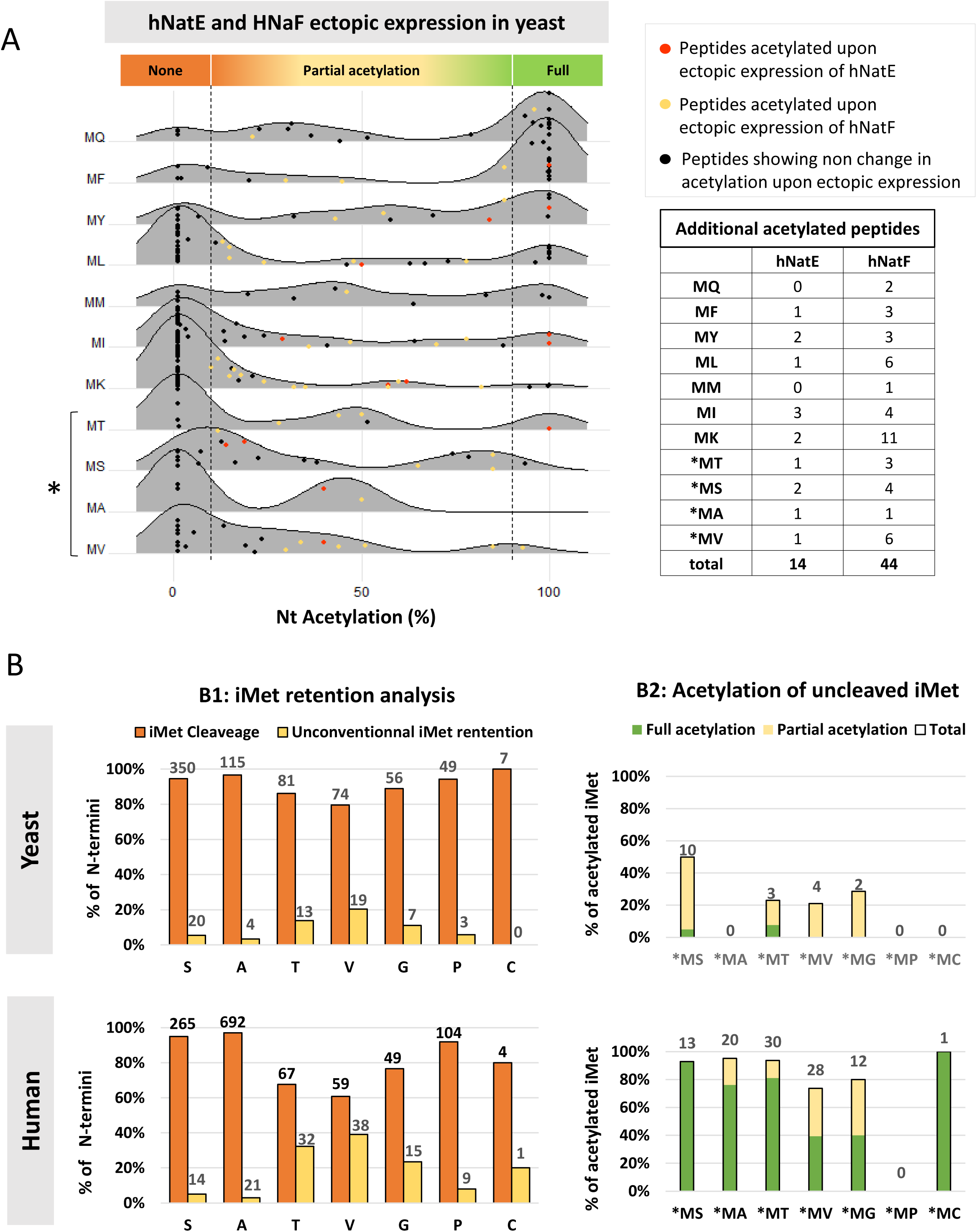
hNatE and hNatF extend iMet acetylation and compete with iMet removal Ectopic expression of hNatE and hNatF in yeast (A) generates new acetylated peptides, shown as colored dots (orange: hNatE; yellow: hNatF; black: no change) on ridgeline plots (left: control yeast Nt-acetylome; right: upon hNat expression) with peptides classified by their N-terminal residues. All additional Nt-acetylation, summarized in the right-hand table, targets the initiator methionine, including iMets normally removed by MAPs (*), reflecting a competitive effect. Analysis of N-terminal ends with unconventional iMet retention (B), although followed by small residues (S, A, T, V, G, P, C) typically targeted by MAPs, revealed higher iMet retention in humans compared with yeast (B1: dark and light orange bars, corresponding respectively to iMet removal and retention rates), associated with increased acetylation levels (B2: green and yellow bars, corresponding respectively to partial and full acetylation rates). Overall, these data indicate that hNatE and hNatF likely contribute to higher rates of iMet acetylation in humans compared to yeast by acetylating NatB/NatC-type iMets and promoting the formation and acetylation of atypical N-termini through competition with MAPs.

Interestingly, the frequencies of unusual iMet retention vary depending on the N-terminal residue and species. Among the unconventional N-termini resulting from unusual iMet retention, MT-, MG-, and MV-types are the most frequently observed, and are much more prevalent in humans than in yeast (Figure 3B, left panel, Table S1). Moreover, iMet NTA of these unconventional N-termini is rare and partial in yeast but nearly systematic in humans (Figure 3B, right panel). These quantitative analyses strongly support the previously proposed model of competition between MetAPs and Nats^6,8,9^, which increases the diversity of possible Nt-proteoforms (with or without iMet, acetylated or non-acetylated) in humans. Recent cryo-EM structures of ribosomes revealed that NatA and MetAP bind simultaneously for efficient processing of NatA type substrates^15^. Our quantitative analysis suggests that the MT-, MG-, MV-type N-termini are less well recognized by the MetAP/NatA enzyme pair. This opens the way to possible competition for cotranslational or post-translational NTA of unconventional iMets, particularly in clades such as metazoans expressing three different Nats capable of this activity (NatC/E/F)^1^. Owing to the limited coverage of proteomes by COFRADIC data, the identification of unconventional iMet N-termini is restricted to approximately one hundred instances (66 in yeast and 131 in humans). Consequently, this area of study remains largely unexplored and presents significant challenges for researchers. More generally, our analysis demonstrated that many Nats substrates are still unknown and difficult to predict. It highlighted the fact that escape from MetAP and/or NTA activity widens the range of possible Nt-proteoforms, and underscores the need to develop other, more sensitive approaches to detect Nats targets *in vivo*.

### Global analysis of mRNAs co-immunoprecipitated with Nats expanded their set of potential substrates and revealed a cryptic group of targets produced by alternative translation starts

To overcome sensitivity limitation of the Nt proteomic approach, we have applied in yeast the Selective Affinity Purification of Translating Ribosomes (sel-TRAP) method, to capture by immunoprecipitation NatA, NatB and NatC associated nascent chains, as well as their corresponding mRNAs, which we characterized using transcriptomic analyses^14^ (Figure 4A, Table S2). This approach proved highly effective in detecting the cotranslational interaction of NatC with mitochondrial precursors^14^. The latter are in very low quantity in cells and virtually undetectable with Nt proteomics, due to their rapid import into mitochondria and subsequent cleavage of their Nt Mitochondrial Targeting Sequence (MTS). Here, we have analyzed these data in more details to complete our knowledge of the putative substrates of NatA, NatB and NatC and propose the most comprehensive list of cellular targets for each Nats based on experimental data.

**Fig 4.**
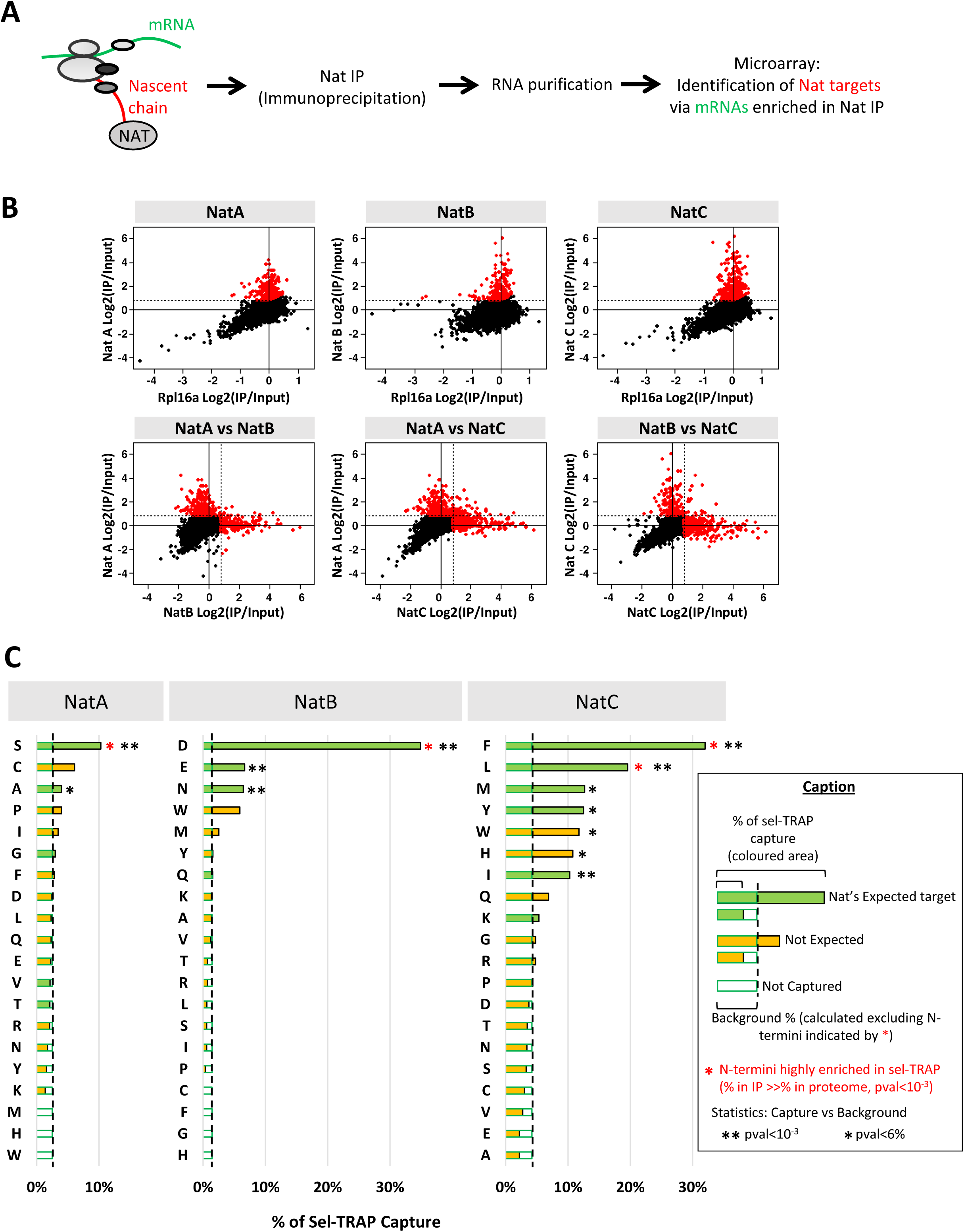
Co-translational capture of NatA, NatB, and NatC targets by sel-TRAP sel-TRAP strategy (A) was used to detect co-translational targets of Nats in yeast. NatA, NatB, and NatC were immunoprecipitated from cells expressing Ard1-PA, Nat3-PA, and Mak3-PA, capturing ribosomes with Nat-associated nascent chains. Enrichments of mRNAs (B) in IP versus total extracts (log₂ IP/Input) were measured by microarray, and putative targets of Nats (red dots in B) were defined as those with enrichment > 0.8 and significantly higher than Rpl16A-PA control IP, as shown in dot plots comparing each Nats IP to the non-specific ribosome control pool (B, upper panels). Comparison of enrichment values across the three Nat IPs confirmed minimal overlap among their respective sets of targets (B, lower panels). Capture efficiency analysis (C) for each N-terminal type in the NatA, NatB, and NatC target lists confirmed expected substrate specificities and revealed unanticipated targets. Bar plots show the recovery percentage for each N-terminal type, defined as the fraction of proteins with this residue recovered in the target list. Expected residues (based on COFRADIC specificity, Fig. 2) are shown in green, others in yellow. Residues significantly enriched in each IP (i.e., overrepresented in the target list compared with their frequency in proteome; p < 10⁻³) are marked with red stars. Remaining residues were used to estimate the mean background capture level (dotted line), representing maximal contamination (2.4% NatA; 1.4% NatB; 4.1% NatC). N-termini captured above this threshold were considered significantly retained (** p < 10⁻³; * p < 6%).

We identified a set of mRNAs significantly enriched in the Nats sel-TRAP assay relative to the control experiment, which consisted of non-selective ribosome immunoprecipitation using RPL16a as bait. The resulting putative target sets (enrichment > 0.8 and significantly higher than in the Rpl16a IP) comprised 244 candidates for NatA, 180 for NatB, and 336 for NatC (red dots in Figure 4B), with minimal overlap among them, consistent with the known substrate-specific selectivity of individual Nats (Figure 4B). To evaluate the specificity of this approach, we examined the distribution of amino acids at position 2 (immediately following the initiator methionine) within these candidate lists (Table S2).

First, statistical analysis revealed a significant enrichment (hypergeometric test, p < 10⁻³) of N-termini starting with serine in NatA targets (59% vs 25% in the proteome), aspartate in NatB targets (59% vs 5%), and leucine and phenylalanine in NatC targets (17% and 20% vs 3% and 6%), consistent with expected substrates of the respective Nats (Figure 2). However, this data-analysis strategy lacks sensitivity for detecting substrates that are recognized inconsistently by Nats or are rare in the yeast proteome (Figure 1A, peptides counts), as these may escape enrichment analysis.

To address this limitation, we applied a complementary approach. sel-TRAP recovery rate was calculated for each N-terminal type (fraction of N-termini in the sel-TRAP lists relative to their total occurrences) and ranked in descending order (Figure 3C, % sel-TRAP capture). We then applied a statistical approach to distinguish specific capture from background. Background capture (Figure 3C, dotted lines) was estimated as the fraction of N-termini recovered in the sel-TRAP lists after excluding those previously identified in the enrichment analysis (Figure 3C, red stars). Each N-terminal category was subsequently tested against this background using a binomial test, and categories with capture values significantly above background (p < 10⁻³ or p<0,06, see below) were defined as specifically captured (Figure 3C, black stars).

The identified N-termini corresponded to the expected substrates of the respective Nats (green bars, Figure 4C), consistent with COFRADIC validation (Figure 2). For NatB, most of the expected substrate types were recovered, except for MQ- N-termini, which also showed lower Nt-acetylation in the COFRADIC data (Figure 4C). NatA sel-TRAP generally exhibited low capture efficiency, with the notable exception of S- N-termini (Figure 4C), which were consistently more efficiently Nt-acetylated than other NatA-type substrates according to COFRADIC data (Figure 2A1). For NatC, the initial stringent cut-off (p < 10^−3^) detected only MF-, ML-, and MI-type N-termini, whereas a less stringent criterion (p < 0.06) allowed recovery of all typical substrates (Figure 4C, single black stars). Notably, this analysis also identified MH- and MW- as putative new NatC substrates, previously undetected in the COFRADIC data (Figure 4C).

To clarify whether the unexpected sel-TRAP candidates (N-termini not matching COFRADIC data; yellow bars, Figure 4C) represented background contaminants or new Nat substrates, we examined their representation in the target lists based on increasing enrichment thresholds (log₂(IP/Input)) applied for list definition (Figure 5A, Table S3). The expected targets (Figure 5A, upper panel) were highly enriched in the lists obtained, particularly for NatA and NatB, representing up to 100% of the top-ranked targets (enrichment > 3), compared to 52% and 18% in the proteome. In contrast, for NatC, expected targets represented only 76% of the top-ranked proteins (compared with 21% in the proteome), as 24% corresponded to non-expected N-termini. This may reflect background contamination or, alternatively, suggest that NatC recognizes additional N-termini not captured by COFRADIC analyses. Previous studies reported that NatC preferentially acetylates N-termini with an arginine at position 3. Consistently, we observed significant enrichment of these N-termini having an arginine at the 3^rd^ position among the unexpected NatC targets with the highest enrichment values (Figure 5A, bottom panel; Figure 5B, Table S3). The top ten unexpected NatC targets (Table S3) include three MHR- and two MQR- N-termini. The remaining five top unexpected NatC targets correspond to MetAP/NatA substrate types (MS-, MA-, MT-, and MG-, Figure 5B), whose recognition by NatC could result from incomplete iMet removal. Alternatively, these unexpected NatC targets could result from alternative translation start sites generating proteoforms with N-termini matching canonical NatC recognition motifs. For example, the mRNA with the highest enrichment in the NatC sel-TRAP dataset (enrichment = 6.1) encodes the mitochondrial protein MRP13. Although its annotated sequence begins with MG-, it contains a typical NatC-specific MFR- motif 27 nucleotides downstream of the annotated iMet. Given NatC’s very high affinity for MRP13, it is tempting to speculate that a truncated form of MRPL13 might be produced via alternative translation start site selection, generating an Alt-start proteoform with strong NatC recognition.

**Fig 5.**
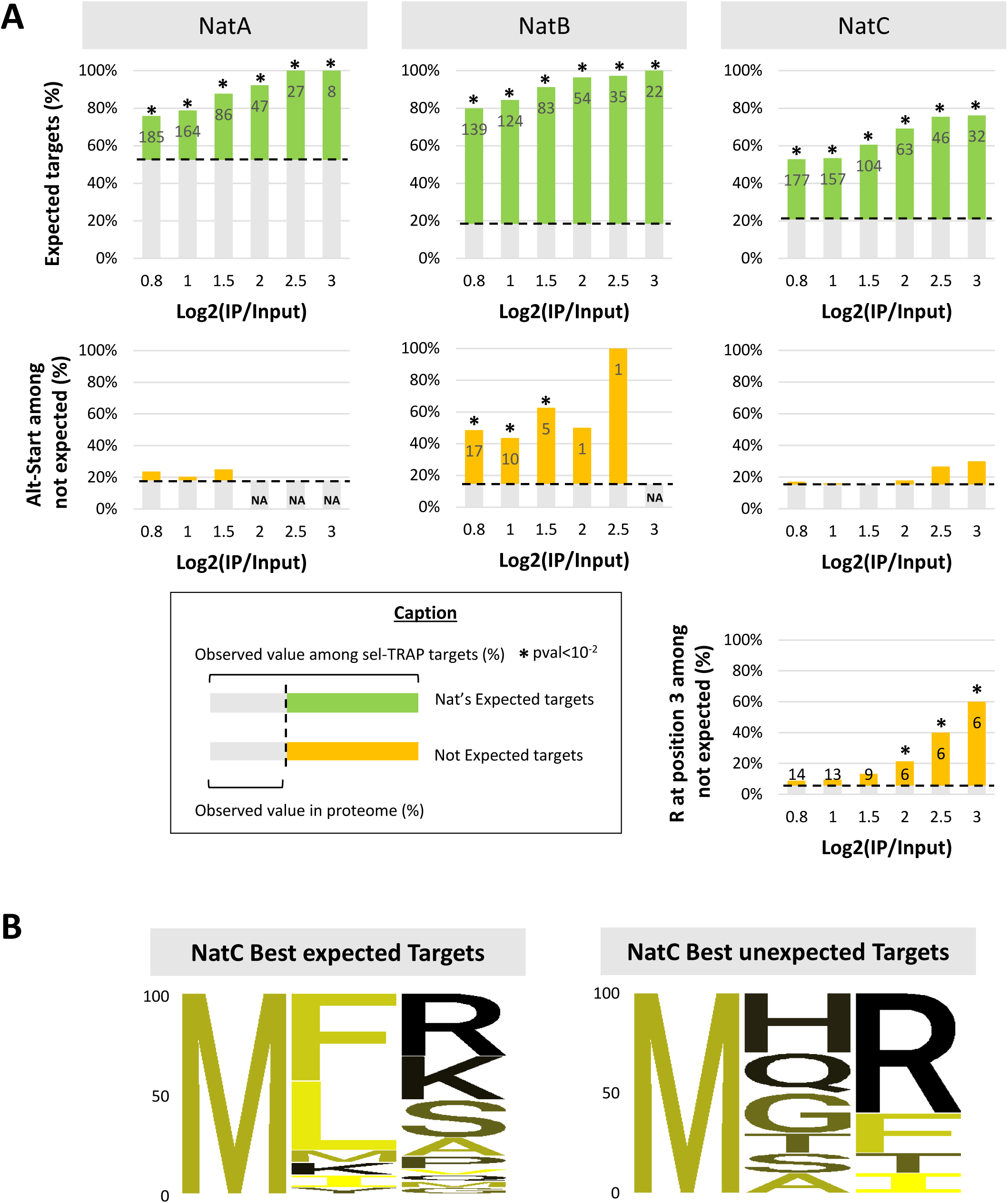
sel-TRAP uncovers novel Nat substrates and alternative N-terminal proteoforms Analysis of sel-TRAP target lists (A) shows overrepresentation of expected substrates (green bars, top panels). Unexpected targets (yellow bars, middle and bottom panels) may result from alternative N-terminal proteoforms with compliant N-termini (enriched Alt-starts in NatB IPs) or, for NatC, from extended specificity when the N-termini contain arginine at position 3 (6 cases among the 10 unexpected targets with the highest scores). Bar graphs show the proportion of each category (top, expected targets; middle, compliant Alt-starts among unexpected targets; bottom, Arg3 among unexpected NatC targets) stratified by enrichment thresholds used to define the target list (0.8–6), with the number of targets in each category indicated. Dotted lines represent proteome frequencies, and stars mark significant enrichment (p < 10⁻²). Only compliant Alt-starts within 50 codons of the annotated iMet were included. N-terminal amino acid analysis (B, sequence logos for positions 1–3) confirmed Arg3 enrichment in NatC top targets, seen both among expected substrates (left, log₂ IP/Input > 6) and in the ten top unexpected targets (right, log₂ IP/Input > 6), which included three MHR- and two MQR-type N-termini.

Based on this observation, we conducted a systematic analysis of potential proximal alternative translation start sites (Alt-starts located up to 50 nucleotides downstream of the annotated iMet) to determine if this phenomenon could account for some of the unexpected targets identified in sel-TRAPs datasets (Figure 5A, middle panel and table S3 and S4). We found 14 potentially compliant Alt-starts among the unexpected targets of NatA, 17 among those of NatB and 27 among those of NatC. These Alt-start candidates are significantly more frequent than expected by chance in the NatB dataset, accounting for 49% of unexpected candidates, whereas NatB-type proximal Alt-starts are found in only 15% of the proteome. Importantly, our observations align with recent yeast ribosome profiling analyses, which revealed hundreds of extended or truncated protein forms from alternative start-site selection^16^. Notably, several of the potential Alt-starts we identified were also reported in that study and in the COFRADIC data (see Table S4 and next Results section). Thus, our analysis of the sel-TRAP data highlights that prediction of Nats targets based solely on analysis of their annotated coding sequence could potentially be further complicated by unknown variations in ribosome selection of start codons. Given that alternative start annotation is still in its infancy, this phenomenon represents an additional challenge for accurately characterizing Nat targets and for appreciating the resulting diversity of cellular Nt-proteoforms.

### Integration of Omics data: identification of canonical proteins targeted by NATs and discovery of cryptic substrates from alternative translation start sites

Extrapolations from Nt proteomic data previously suggested that 60% of the yeast proteome could undergo NTA^1,2^. Yet, our meta-analysis of Nt-COFRADIC data revealed that it is difficult to predict the NTA level of a given protein, particularly for N-termini that are poorly acetylated and/or represented in these data. NTA prediction is indeed complicated by NTA escapes, which include the absence or partial acetylation of some putative Nats substrates, as well as Nt maturation crosstalks, where MetAP and Nats compete for a given substrate, leading to unconventional iMet acetylation. Additionally, our sel-TRAP analysis uncovered cryptic Nats targets potentially explained by Alt-start selection during translation, expanding our NTA inquiry to previously overlooked Nt-proteoforms.

In order to compile from the Omics data a comprehensive list of proteins targeted by NatA, NatB and NatC, including canonical proteins and Alt-start Nt proteoforms, we first revisited the substrate specificities of Nats. Based on our analysis of Nt-COFRADIC and sel-TRAP data, along with literature on *in vitro* activity assays^17–20^ (Table S5), we present an integrated statistical view of NatA, NatB, and NatC substrate preferences (Figure 6A). From the COFRADIC analysis, confidence scores were assigned to each substrate type according to the percentage of Nt acetylated proteins and the number of detected N-termini (see COFRADIC score scale in Table S5). Similarly, we assigned confidence scores from statistical analysis of sel-TRAP data (see sel-TRAP score scale in Table S5). While Nt-COFRADIC provides direct information on the NTA levels of proteins, sel-TRAP, which relies on affinity capture, gives an overview of the affinity of Nats for their substrates. The data obtained for NatA and NatB are very consistent since substrates types with the highest NTA levels detected in COFRADIC also showed better captured in sel-TRAP. They indicate a stronger preference of NatA for S- termini and variable affinities of NatB for its substrates, with high affinity for MD-, intermediate affinity for MN-and ME-, and lower affinity for MQ-. The Nt-COFRADIC data poorly represent the N-termini targeted by NatC, resulting in unreliable predictions of NatC substrates (Figure 2 A2). In contrast, sel-TRAP analysis provided valuable information on NatC substrates. It confirmed NatC *in vivo* preferences for N-termini with large hydrophobic amino acids, and uncovered previously unknown putative NatC substrate, such as N-termini starting with MH and MQ, followed by an arginine in position 3 (Figure 5B). This reinforces the established role of the Arg3 residue in substrate recognition by NatC and suggests that NatC’s substrate selectivity extends beyond large hydrophobic residues in position 2, with position 3 also playing a critical role.

**Fig 6.**
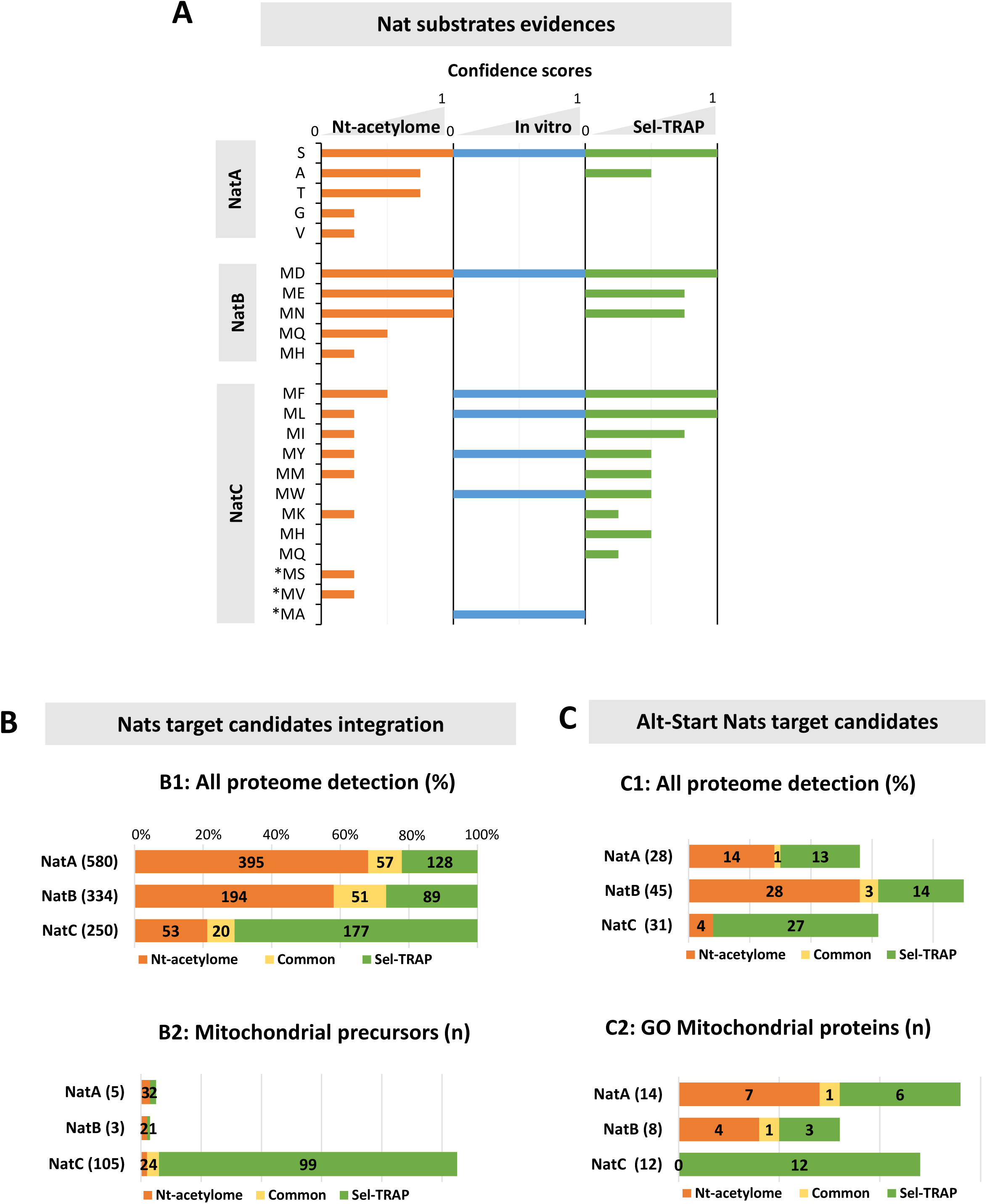
N-terminal data integration: Nats substrate specificities and compilation of canonical and Alt-start target lists Substrate specificities of Nats (A) were inferred from Nt-COFRADIC, *in vitro* assays, and sel-TRAP data. Confidence scores (0, 0.25, 0.75, 1) were assigned based on quantitative and statistical criteria (see Table S5), with *in vitro* scores set to 1 based on literature evidence. COFRADIC scores reflect the number and proportion of acetylated peptides with compliant N-termini, while sel-TRAP scores combine enrichment against proteome frequencies, binomial testing versus background capture (Fig. 4), and analysis of top NatC targets (Fig. 5). Integration of Omic data for Nat target definition (B, C): NatA, NatB, and NatC target lists were compiled (B1, C1) from COFRADIC (Nt-ac >10%) and sel-TRAP (score >0.8) datasets, assigned according to the substrate specificity defined in A. Canonical targets (B) start at the annotated iMet, while Alt-start targets (C) include COFRADIC-detected N-terminal proteoforms or unexpected sel-TRAP targets with an alternative start within 50 codons producing a compliant N-terminus. Analyses of mitochondrial precursors with N-terminal MTS (B2) or GO-annotated mitochondrial proteins (C2) revealed enrichment among NatC targets. Bar graphs show counts of canonical (B1/B2) and Alt-start (C1/C2) targets for each Nat and detection method (orange: COFRADIC, green: sel-TRAP, yellow: both).

From this analysis, we compiled the most comprehensive list of canonical proteins (canonical translation start) identified as Nat substrates in yeast based on experimental data (Figure 6B and Table S4). Nat targets were identified from the Nt-COFRADIC data based on their Nt-acetylation levels (>10%) or from the list of candidates enriched in the sel-TRAP data (enrichment >0.8 and significantly higher than in the Rpl16a IP). Candidates for each Nats were selected according to our integrated substrate selectivity analysis (Figure 6A). In each dataset, we selected target candidates for Nats based on residues in position 2: for NatA, proteins with S, A, T, V, or G; for NatB, those with D, N, E, Q, or H; and for NatC, those with F, L, I, Y, M, W, or K. The list of putative NatC targets identified by sel-TRAP was completed by candidates with an H or Q in position 2, as our analysis of the sel-TRAP data indicated that these NatB substrates could also be recognized by NatC. After excluding 20 sel-TRAP candidates with detected N-terminal acetylation below 10% (as measured by COFRADIC), the final dataset comprises 1,145 Nat target candidates, representing nearly 20% of the yeast proteome. Of these, 770 proteins were identified by our meta-analysis of Nt-COFRADIC data, and their NTA levels are therefore available. Nt proteomic data suggested that 60% of the yeast proteome might undergo NTA^1^, but since this approach only covers highly expressed proteins, the detected Nt-acetylated proteins represent less than 15% of the proteome. sel-TRAP enabled us to expand the list of putative Nat targets. In particular, while our meta-analysis of Nt-COFRADIC identified only 73 NatC substrates, sel-TRAP enhanced our understanding of NatC *in vivo* selectivity by revealing 177 new putative targets, many of which are mitochondrial precursors that are difficult to detect with proteomics (Figure 6B).

sel-TRAP has also provided new insights into Nt-Acetylome dynamics, highlighting the role of alternative translation start sites in generating cryptic Nat substrates. This led us to compile a list of cryptic Nat targets likely resulting from Alt-start translation (Figure 7A, Table S4). While sel-TRAP analysis identified 58 such cases, revisiting the Nt-COFRADIC data revealed 50 Nt-acetylated peptides that may correspond to Nt-proteoforms resulting from translation initiation at a methionine within the first 50 amino acids downstream of the annotated start site (proximal Alt-start). The final list of 104 cryptic Nat substrates detected in Nt-proteomic and/or sel-TRAP data includes 28 for NatA, 45 for NatB, and 31 for NatC (Figure 7A). Nt-COFRADIC and sel-TRAP data are poorly overlapping (Figure 6B), and only four cryptic Nat substrates were detected by both approaches (Figure 7A), most likely because each method captures only a small and distinct fraction of them due to their respective sensitivity limits. Recently, with a modified ribosome profiling approach for translation start site detection, the Brar laboratory identified 388 truncated Nt-proteoforms of annotated proteins in yeast^16^. Among them, 235 were proximal truncations (within 50 aa of the annotated start codon), likely originating from full-length transcripts through alternative start selection. Thirteen of them were also predicted in our analysis (8 predicted in COFRADIC, 3 in sel-TRAP and 2 in both), further supporting our detection of actual cryptic Nat substrates arising from alternative start site selection (see Table S4). This includes an alternative Nt-proteoform of fumarase (Fum1), detected with the ribosome profiling but also by both COFRADIC and sel-TRAP, and which we will discuss further below. Yet, gaps remain in the mapping of Nt-proteoforms, as reflected by the low overlap between different detection approaches, suggesting that, even with ribosome profiling, a large proportion of alternative translation start events remains undetected.

**Fig 7.**
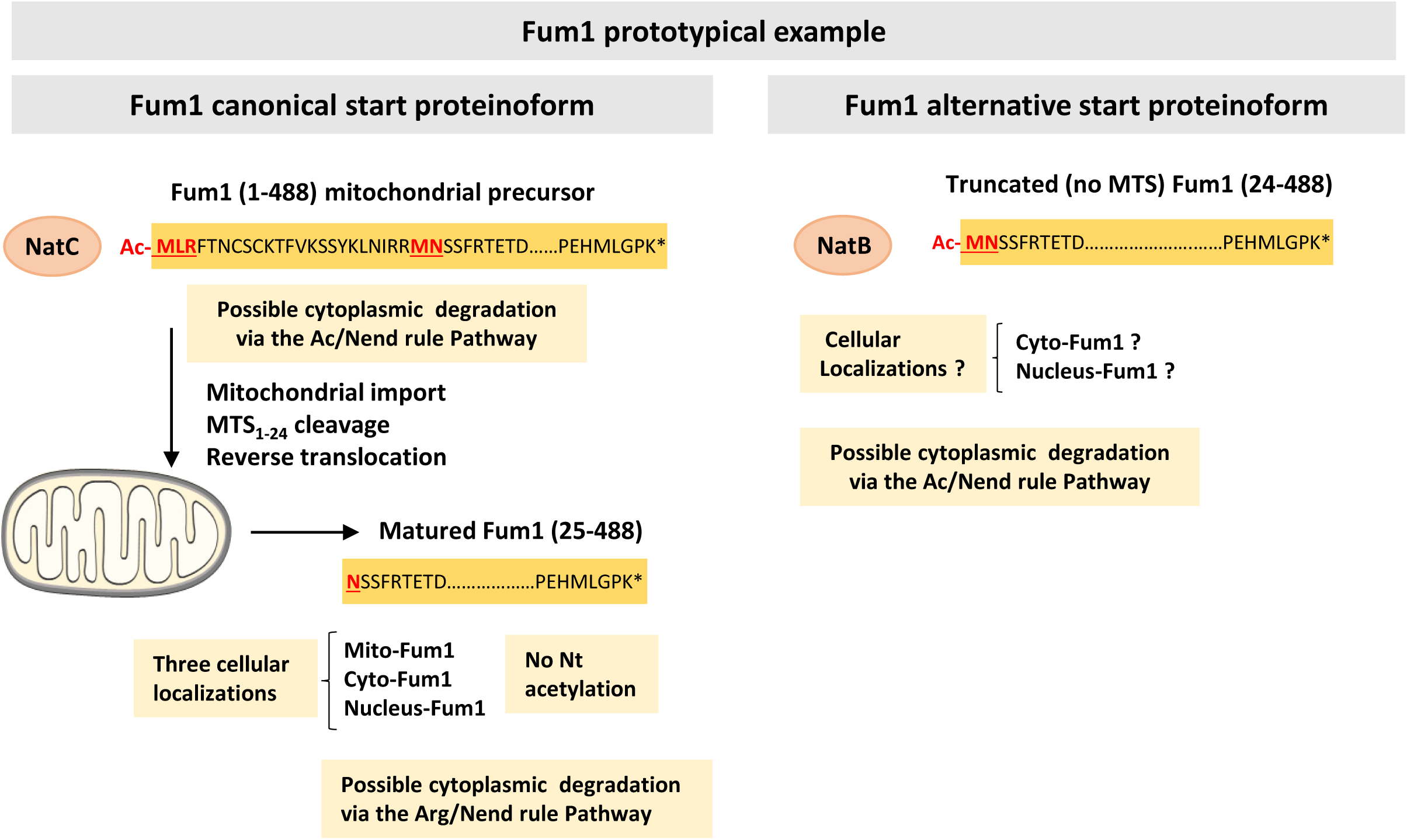
Nt-proteoform dynamics: Fum1 as a prototypical model Fumarase (Fum1) localizes to mitochondria, cytoplasm, and nucleus in yeast and humans. In yeast, canonical translation yields the precursor Fum1(1–488) and the mature mitochondrial form Fum1(25–488) generated by MTS cleavage. A fraction of the latter may be retro-translocated to the cytoplasm^25^. Our data and previous reports^16^ support an alternative start product, Fum1(24–488), lacking the MTS and thus likely not targeted to mitochondria. The three Nt-proteoforms differ in N-terminal acetylation, Fum1(1–488) and Fum1(24–488) being NatC and NatB substrates, respectively, while Fum1(25–488) remains unacetylated, suggesting distinct stability and degradation control through the N-end rule pathway (see main text).

In their ribosome profiling study, the Brar laboratory found that many truncated proteins generated from alternative translation initiation sites display distinct subcellular localizations compared to their annotated full-length counterparts, highlighting a widespread mechanism for dual protein localization^16^. Notably, a common pattern was the loss of the N-terminal mitochondrial localization sequence found in the full-length proteins, leading to their cytoplasmic localization. Similarly, we observed that mitochondrial localizations are enriched among our cryptic Nat substrates, representing 46% and 40% of the NatA and NatC Alt-start substrates, respectively. Among the 104 Alt-start Nt proteoforms we detected, 33 are classified as mitochondrial proteins according to Gene Ontology, with 29 further confirmed in the SGD or UniProt databases (Figure 6C2 and Table S4). Of these, 20 have an N-terminal MTS (either annotated in UniProt^21^ or predicted with MitoFates^22^) which is fully or partially truncated in the Alt-start Nt proteoform. Interestingly, 17 of the 29 mitochondrial proteins in our Nt proteoform list were also observed in other compartments and are classified as dual-localized proteins according to the literature^23^. For five of these proteins (CDC9, GLR1, GRS1, YSA1, and MRX2), the truncated forms responsible for their dual localization have already been described in the literature and correspond to the Nt-proteoforms we detected (see references in Table S4). Brar’s ribosome profiling data^16^ also support the truncated form of GLR1 and five additional Nt-proteoforms of mitochondrial proteins identified in our analysis (GUT1, RDL1, FUM1, ERV1, ETR1, and MRP20). These findings indicate potential mechanisms for dual localization of GUT1, RDL1, and FUM1^23^, and suggests MTS truncated variants for ETR1, and MRP20. For the latter two, no secondary localization has been reported, and in the case of MRP20, the MTS truncation appears partial, with only 10 of the 45 MTS residues removed. Among proteins with exclusive mitochondrial localization, we detected several additional cases of partial MTS truncation (IDP1, MIP1, MRP13, MRPL16, MRP20, MRPL3, RIP1, SDH3) in our sel-TRAP data, most of which are predicted to be cryptic NatC substrates (Table S4). The mitochondrial ribosomal protein MRP13 exemplifies this phenomenon. Although MRP13 shows the highest enrichment in the NatC sel-TRAP dataset, its annotated proteoform carries an MG- N-terminus, typical of NatA targets. However, initiation at methionine 28 would generate an alternative proteoform with an MFR- N-terminus, characteristic of NatC substrates and typical of yeast MTS^14^.

Further investigations are required to validate the entire set of N-terminally truncated proteoforms suggested by our analysis of COFRADIC and sel-TRAP data. Nonetheless, these analyses highlight the role of alternative start site selection in diversifying Nat substrates, thereby expanding the range of Nt-proteoforms (acetylated versus non-acetylated) with potentially distinct functions and/or localizations.

### Discussion on NTA and Nt Proteoform Dynamics: Mitochondrial Precursors as a New Model

N-terminal acetylation modulates protein function and fate by adding a bulky hydrophobic group, which affects folding, interactions, localization, and stability^1^. We have identified new NAT substrates derived from proteoforms with variable N-termini, suggesting that NTA may further diversify the functional repertoire of a given protein. Fumarase (Fum1), a dual-localized mitochondrial protein conserved from yeast to human^24^, provides a relevant prototypical example to discuss these functional dynamics (Figure 7).

Based on the convergent results from sel-TRAP, COFRADIC, and ribosome profiling, our analysis provides robust experimental evidence in yeast for the existence of a truncated Fum1 translation product initiated at a methionine 24 codons downstream of the annotated start site. This truncated form, Fum1(24-488), lacks the mitochondrial targeting sequence (amino acids 1–24 of canonical Fum1) and may account for its cytoplasmic localization. However, the current model of dual localization of yeast Fum1 is based on a single translation product^25^, Fum1(1–488) (Figure 7, left panel), because cytoplasmic localization is still observed in Fum1 mutants lacking Met24. In this model, the cytoplasmic protein Fum1(25-488) is proposed to result from the retrograde movement of the partially imported precursor Fum1(1-488), blocked in a transport-incompetent state within the translation pore, after cleavage of its MTS. Thus, in yeast, three Nt-proteoforms of Fum1 may coexist (Figure 7): the mitochondrial precursor Fum1(1-488), the mitochondrially processed Fum1(25-488) and finally the Alt-start truncated Fum1(24-488). These Nt-proteoforms appear to be differentially targeted by Nats and may display distinct subcellular localizations, which could consequently modulate their interaction networks and overall stability (Figure 7).

Fum1(1–488) precursor exhibits the typical features of a NatC substrate (MLR N-terminus; Figure 7), consistent with the strong enrichment of FUM1 mRNA in our NatC sel-TRAP dataset (enrichment value: 2.92). The mature form, Fum1(25–488), generated by MTS cleavage in mitochondria, is thought to distribute between mitochondria and the cytosol via reverse translocation. Moreover, this cytosolic Fum1(25–488) can translocate into the nucleus in response to DNA damage, where its activity is required for efficient repair of double-strand breaks^26^. This proteoform, bearing an N-terminal asparagine, is not a substrate for any Nat (Figure 7). Finally, the newly identified Alt-start truncated Fum1(24–488) represents a typical NatB substrate, characterized by an MN N-terminus (Figure 7B). It is detected as fully acetylated in Nt-COFRADIC analysis, in agreement with the enrichment of FUM1 mRNA (enrichment value: 1.75) in NatB sel-TRAP data (Table S4). Because this N-terminal proteoform lacks the MTS (amino acids 1–24), it is likely not imported into mitochondria, supporting the notion that two N-terminal Fum1 proteoforms, Fum1(25–488) and Fum1(24–488), coexist in extra-mitochondrial compartments. These forms differ only by the presence of an acetylated N-terminal methionine, specific to the truncated Alt-start variant. This observation raises important questions regarding their respective functions and subcellular localization. Is the Alt-start form also imported into the nucleus, and if so, does its acetylation by NatB contribute to the role of Fum1 in the DNA damage response?

In addition to different cellular localizations and functions, the Fum1(25–488) and Fum(24-488) may also have different stability levels. The N-end rule pathway, part of the ubiquitin-proteasome system, includes branches such as the Ac/N- and Arg/N-end rules, which recognize specific N-terminal residues and their modifications and could potentially direct the diverse Fum1 N-terminal proteoforms to proteasomal degradation^27–29^(Figure 7). However, predicting protein stability solely from the Nt residue is challenging, as engagement in the N-end rule pathways is modulated by protein folding and Nt accessibility^29^. Experimental characterization, including stability assays, of Fum1 N-terminal proteoforms across different cellular compartments (cytosol, nucleus, mitochondria) would provide critical insights into the complex interplay between Nt proteoforms and Nt acetylation in regulating protein function and life cycle.

### Concluding remarks: from Yeast to human Nt-acetylome, a challenge for the future

Our comprehensive exploration of Nt-proteomic data, combined with the newly developed sel-TRAP approach for *in vivo* capture of Nat targets, has provided an integrated view of the Nt-acetylome in yeast and offered new insights into Nat substrate specificities. Nevertheless, our understanding of the cellular N-terminal acetylome remains incomplete. Notably, our analysis revealed an additional layer of complexity in the dynamics of N-terminal proteoforms arising from alternative translation start site selection. This previously overlooked aspect of the N-terminal acetylome likely plays a key role in diversifying protein function, localization, and stability within the cellular proteome.

This issue is particularly relevant in the human context, where the diversity of N-terminal isoforms, resulting from alternative splicing, transcription start site selection, and alternative translation initiation, is further amplified by a range of N-terminal modifications, including the dynamic interplay between methionine aminopeptidases and various N-terminal acetyltransferases (NatC/E/F). This results in a complex landscape of N-terminal proteoforms whose integration into cellular physiology remains largely unexplored^30^. Importantly, both N-terminal sequence variation and alterations in N-terminal acetylation have been implicated in the pathogenesis of various diseases, including neurodegenerative disorders, cancer, metabolic conditions such as type 2 diabetes, and developmental syndromes^30^.

Underscoring the need for further N-terminal profiling in humans, less than 10% of the ∼20,000 canonical protein isoforms annotated in UniProt have been studied by N-terminal proteomics to date (Figure 1B), revealing 1,683 N-terminally acetylated proteins (Table S6), including 991, 470 and 119 attributed to NatA, NatB, and NatC, respectively, and 103 corresponding to atypical iMet acetylation by either NatC, NatE, or NatF. These data not only capture a limited fraction of the cellular N-terminal acetylome but also fail to account for Nt-acetylation of alternative human isoforms. Extending our analysis of Alt-start events from yeast to humans, we identified 123 N-terminally acetylated peptides likely arising from alternative translation initiation within 50 residues of the canonical start codon (Table S6). Detected within only ∼10% of the human proteome, these findings reveal a major blind spot and highlight the need for advanced experimental approaches to systematically capture the full diversity of human N-terminal proteoforms. Meeting this challenge will be essential to achieve a comprehensive and functionally relevant map of the human acetylome.

## METHODS

All analyses were performed using R (version 4.2.2).

### N-terminal Proteomics Data

Yeast and human proteomics datasets generated using COFRADIC^31^ were collected from previously published studies^4–11^ (Table 1 and Sup1 input data). The datasets report peptide sequence, protein information (standard name, UniProt ID, annotation), peptide coordinates, and N-terminal acetylation percentages in wild-type cells and in conditions with altered NAT activity. NAT perturbations were achieved by gene deletion in *S. cerevisiae*, RNA interference or CRISPR knockout in human cells, or ectopic expression of human NATs in yeast.

First, for each dataset, all N-terminal peptides, whether acetylated or not, starting at position 1 or 2 of the corresponding proteins, depending on initiator methionine retention or cleavage, were retained (Table S1). These peptides correspond to conventional protein annotations.

Additionally, we searched the datasets for acetylated peptides potentially arising from proximal alternative translation initiation. Specifically, we retrieved peptides whose N-termini originated from a methionine located within the first 50 amino acids downstream of the annotated start site, whether this methionine was retained (yielding peptides beginning with Met) or cleaved by MetAPs (yielding peptides beginning with the small residues S, A, T, V, G, C or P). Following compilation across datasets, these two classes of peptides comprised 104 in yeast and 123 in human and were designated as Alt-start NAT substrates (Table S4 and S7).

### Co-translational Targets Data

sel-TRAP data^14^ for yeast NatA, NatB, and NatC are publicly available on ArrayExpress (E-MTAB-11772) with detailed protocols. Protein A-tagged NAT catalytic subunits and the ribosomal protein Rpl16a were immunoprecipitated, and co-purifying mRNAs were quantified by microarray using input RNA as reference. Proteins undergoing co-translational acetylation by each NAT were inferred from the mean Log₂(IP/Input) of their corresponding mRNAs across two replicates, with significance assessed against the Rpl16a control using the R Limma package (Log₂ (IP/Input)>0,8 and adjusted p-value < 0.05). Table S2 lists 5,618 UniProt entries with their sel-TRAP values and classification as NatA, NatB, or NatC putative targets.

### COFRADIC Identification of Nats Specificity

COFRADIC datasets generated under conditions of altered NAT activity were used to define NAT substrate specificity. For this analysis, only peptides with N-terminal acetylation (NTA) values available in both the wild-type and the corresponding NAT-perturbed condition within the same dataset were retained. Peptides were categorized as non-acetylated (NTA < 10%), partially acetylated (10% ≤ NTA ≤ 90%), or fully acetylated (NTA > 90%).

To minimize false-positive detection, only peptides detected as acetylated in at least one of the two compared conditions (wild-type vs. NAT perturbation) were included in the analysis. Peptides exhibiting significant changes in NTA values upon NAT depletion or ectopic NAT expression were identified based on an absolute log₂ acetylation ratio ≥ 2. A default NTA value of 1% was assigned to peptides lacking detectable acetylation to allow ratio calculation.

NAT substrate specificity was determined from the analysis of N-terminal residues (iMet if retained and position 2) of these peptides. For yeast, substrate assignment was based on datasets from strains lacking the catalytic subunits of NatA, NatB, or NatC (Supplementary Figure 1). For human, substrate specificity was first inferred from ectopic expression of hNatA–F in yeast and subsequently confirmed and expanded using datasets in which NAT activity was perturbed directly in human cells (Supplementary Figure 2 for hNatA–C and Figure 3A for hNatE–F).

### COFRADIC data integration for meta-analysis Nt-acetylome

Wild-type N-terminal acetylomes corresponding to canonical peptides were combined, resulting in datasets of 1,172 and 1,931 entries for yeast and human, respectively (Table S1). For peptides present in multiple datasets (880 in yeast and 1,423 in human), acetylation percentages were averaged, and peptides were retained only if values differed by less than 20% (8 peptides excluded in yeast and 15 in human). For yeast dataset 5 (Table 1), N-terminal acetylation values from the total protein extract were used when available; otherwise, values were taken from the cellular fraction showing the maximal difference between wild-type and NatC-depleted cells, to maximize sensitivity for detecting NatC targets.

Acetylation distributions within each peptide class, defined according to their N-terminal residue and NAT-substrate type (see below), were visualized as ridgeline plots, with density curves representing the overall distribution of peptide acetylation and individual values shown as dots. The proportion of acetylated peptides, including both fully and partially acetylated species, was then calculated, with 5% confidence intervals obtained using the exact binomial test in R.

Across the combined datasets, peptides were further categorized based on the presence or absence of the initiator methionine (iMet). A total of 732 yeast peptides and 1,240 human peptides starting at position 2 were annotated as iMet-cleaved; all matched the canonical MetAP substrate specificity for small residues following iMet (S, A, T, V, G, C, P). Peptides starting at position 1 (iMet-retained) were flagged as exhibiting unconventional iMet retention when the retained iMet was followed by one of these small amino acids (66 peptides in yeast and 131 in human).

Conventional peptides starting at positions 1 or 2 (i.e., excluding unconventional iMet retention) were used to classify the corresponding proteins as NatA, NatB, or NatC substrates, based on Nats specificity defined from COFRADIC analyses of cells with NAT perturbations. Similarly, acetylated peptides from the Alt-start list were classified as NatA, NatB, or NatC substrates.

Based on analyses of NatC depletion and ectopic expression of hNatE or hNatF in yeast, N-terminal acetylation of peptides with unconventional iMet retention was attributed to NatC in yeast or NatC/E/F in human.

These data were used to compile the list of Nats substrates identified from the COFRADIC analyses (Table S4 and S7).

### sel-TRAP analysis of Nats specificity

NAT specificity from sel-TRAP data was determined by analyzing the residue following the canonical iMet in lists of putative targets selected as previously described (Log₂ (IP/Input)>0,8 and Limma adjusted p-value < 0.05). In this analysis, protein N-termini sharing the same first 10 residues were counted only once.

First, hypergeometric tests were performed, as described previously^14^, to identify N-termini significantly enriched in the target lists compared to their representation in the proteome (p-value < 10^−3^). To increase sensitivity in detecting NAT specificity, background capture was estimated as the percentage of N-termini recovered (number of N-termini in the sel-TRAP target list divided by their total occurrences in the proteome), excluding those highly enriched identified in the initial enrichment analysis. A binomial test in R was then applied to identify N-termini with recovery values significantly exceeding the background (p-value < 10^−3^ or < 6%). N-terminal residues identified by this approach were generally compliant with Nat substrate specificity defined by COFRADIC analysis, except for MW- and MH-type residues, which were identified as potential additional NatC substrates.

N-termini in sel-TRAP target lists were classified as expected targets when the residue following iMet matched the corresponding NAT substrate specificity defined by COFRADIC analysis; otherwise, they were classified as unexpected targets. Alt-starts located within the first 50 residues downstream of the canonical iMet were then retrieved from the unexpected target group, and those whose N-terminal residue matched the corresponding NAT substrate specificity were defined as compliant Alt-starts. The proportions of expected, unexpected, and compliant Alt-start targets were analyzed across lists generated with increasing enrichment thresholds (0.8–6). Enrichment relative to proteome frequencies was evaluated using a hypergeometric test (p < 10⁻²). For the NatC unexpected targets, the distribution of those with an arginine at position 3 was additionally analyzed across increasing enrichment thresholds, which allowed the identification of MH- and MQ-type N-termini as potential new NatC substrates (see Results section).

From sel-TRAP data, lists of canonical proteins were compiled for NatA, NatB, and NatC based on N-termini compliant with their respective NAT substrate specificity (as defined by COFRADIC), and by including additional NatC targets with MW-, MH-, and MQ-type N-termini. Targets with detected N-terminal acetylation below 10% in COFRADIC data were excluded from these lists. Lists of Alt-start targets were generated from the initial lists by selecting unexpected targets that contained compliant Alt-starts.

### NAT Substrate Specificity Confident Scores

Three confidence scores were assigned to each NAT substrate type, reflecting evidence from COFRADIC, sel-TRAP, and published in vitro studies (Table S5).

COFRADIC scores were assigned based on the acetylation percentage of each N-terminal type (including fully and partially acetylated peptides) and the number of peptides with available data:

- Score = 1: acetylation > 90% and peptide number > 40
- Score = 0.75: acetylation > 45% and peptide number > 40
- Score = 0.5: acetylation > 67% and 10 ≤ peptide number ≤ 40
- Score = 0.25: acetylation < 67% or peptide number < 10

sel-TRAP scores were assigned based on two criteria: the enrichment p-value from the hypergeometric test and the p-value from the binomial test comparing N-termini recovery to background capture. For the lowest score (0.25), NatC targets with the highest enrichment levels (log₂[IP/Input] > 3) were specifically considered (Results and Fig. 5). Scores were assigned as follows:

- Score = 1.00: p-value (hypergeometric) < 10^−3^
- Score = 0.75: p-value (hypergeometric) ≥ 10^−3^ & p-value (binomial) < 10^−3^
- Score = 0.50: p-value (hypergeometric) ≥ 10^−3^ & 10^−3^ < p-value (binomial) < 6%
- Score = 0.25: NatC best candidate analysis (see Results)

*In vitro* scores were assigned based on published evidence of activity of NatA^17,18^, NatB^19^ or NatC^20^ on peptides with the corresponding N-terminal type. Score = 1 if at least one study reported activity; score = 0 otherwise.

### Analysis of Mitochondrial Proteins

Yeast mitochondrial proteins were identified using Gene Ontology annotations corresponding to the mitochondrion term (GO:0005739) from the GO Consortium database^32,33^. Mitochondrial localization was verified using UniProt^21^ and/or the Saccharomyces Genome Database^34^. Proteins annotated as exclusively mitochondrial or dual-localized were retrieved from the mitochondrial protein list provided in Table S4 of Pines et al^23^. N-terminal mitochondrial targeting sequences were obtained from UniProt^21^ or predicted using MitoFates^22^.

## SUPPLEMENTARY INFORMATION

The supplementary information refers to additional data, and includes several tables.

COFRADIC input data are grouped in a single archive referred **supplemental file S1.**

**Tables S1-S7** are available as independent excel files as described below:

**Table S1:** Combined COFRADIC data for yeast and human and related analysis (proteome coverage, percentage of acetylation and iMet retention)

**Table S2:** sel-TRAP data and position 2 analysis (Enrichment and capture % analysis)

**Table S3:** sel-TRAP unexpected substrates analysis

**Table S4:** Yeast canonical and Alt-start Nats Substrates Integration from COFRADIC and sel-TRAP data

**Table S5:** Scores definition for Nats substrates evidences (COFRADIC, sel-TRAP, *In vitro*)

**Table S6:** Human canonical and Alt-start Nats Substrates Integration from COFRADIC

**Table S7:** COFRADIC data following Nat perturbation in *S. cerevisiae* or human cell lines

**Supplementary Figures S1 and S2** recapitulate the results of Nat perturbation, which allowed the identification of NatA, NatB, and NatC substrate types.

## Supplementary Figures Legends

**Figure S1.**
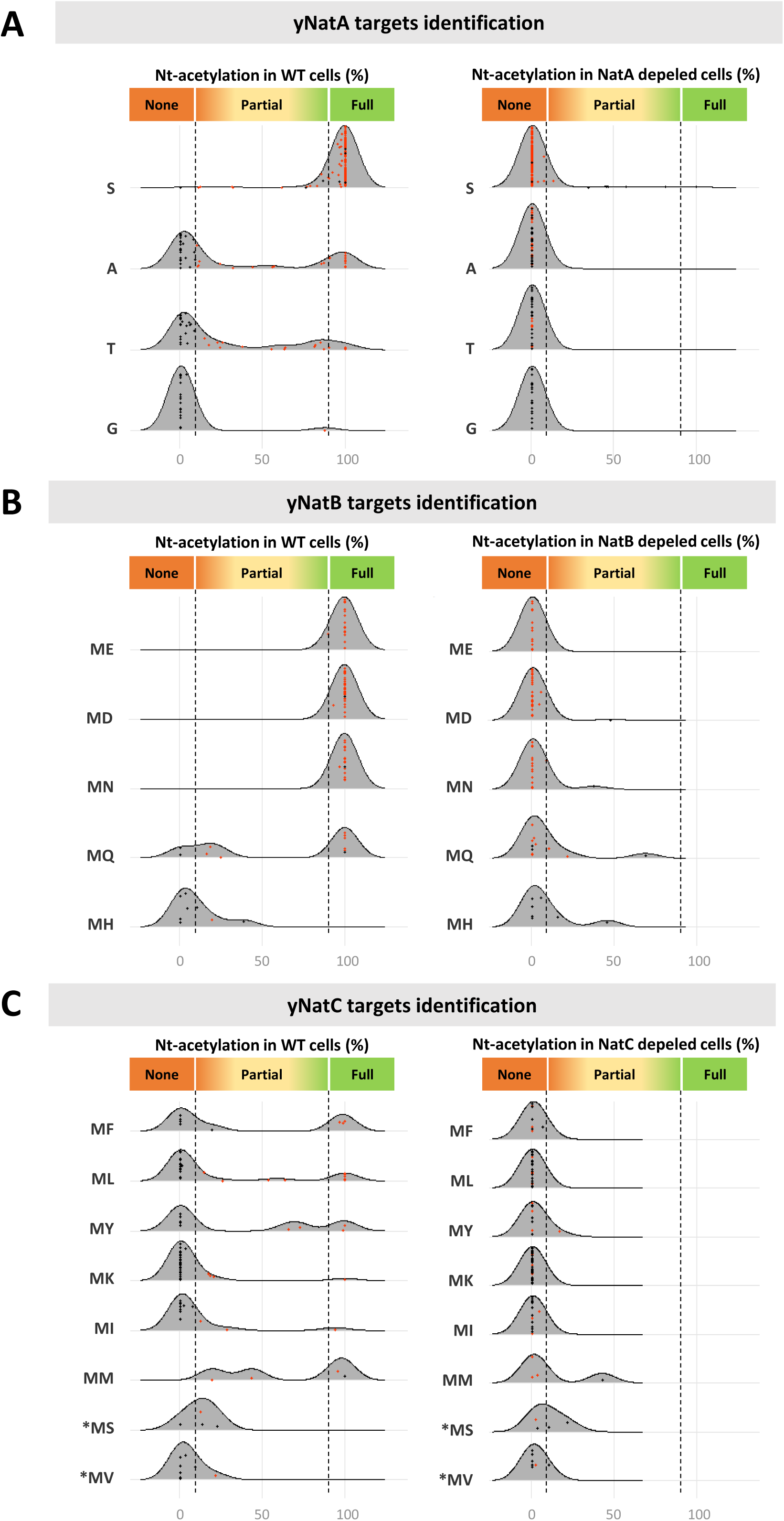
Nat depletions in yeast reveal NatA, NatB, and NatC substrate specificities Depletion of NatA (A), NatB (B), or NatC (C) in yeast causes loss of Nt-acetylation, shown as colored dots on ridgeline plots (red: loss of acetylation; black: no change) comparing control yeast Nt-acetylome (left) and Nat deletion strains (right). Peptides are classified by their N-terminal residues; details on peptides with lost acetylation are in Table S7. These data were used to define Nat substrate specificities applied for peptide classification in the COFRADIC data meta-analysis (Figure 2A). Data used for these analyses were obtained from published dataset 2-5 (see Table1).

**Figure S2.**
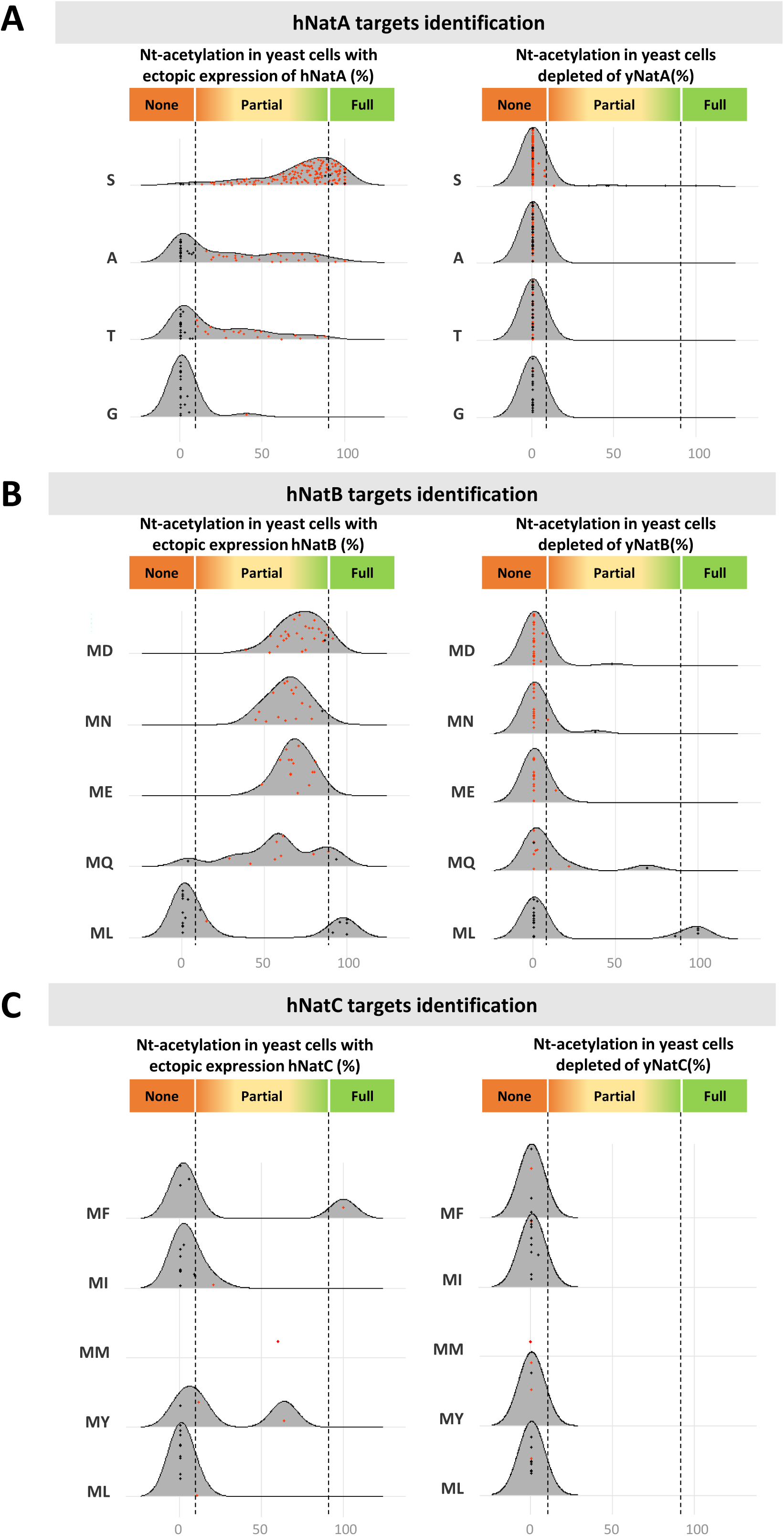
Ectopic expression of hNatA, hNatB, and hNatC in Nat-depleted yeast strains reveals hNat substrate specificities Ectopic expression of hNatA (A), hNatB (B), or hNatC (C) in the corresponding Nat-depleted yeast strains generates new acetylated peptides, shown as colored dots on ridgeline plots (red: gain of acetylation; black: no change), comparing hNat-expressing Nt-acetylomes (left) to Nat-deletion controls (right). Peptides are classified by their N-terminal residues; details on peptides with gain of acetylation are in Table S7. These data define hNat substrate specificities for peptide classification in the COFRADIC meta-analysis (Figure 2B), using datasets 2, 3, and 5 (Table 1). Additional human cell line datasets (7–10, Table 1) revealed specificities not detected in yeast: MV for hNatA, and ML and MK for hNatC (see Table S7). hNatB specificity for ML was not retained, as it relied on a single peptide (see the unique red dot in the hNatB panel for ML N-termini).

## Notes

### Competing Interest Statement

The authors have declared no competing interest.

### Summary of Updates

Figures were inadvertently omitted in the previous version and have now been included.

### Funding

- Sorbonne Université
- Fondation ARC pour la Recherche sur le Cancer: PJA 20171206624
- Centre National de la Recherche Scientifique

